# Tension-Induced Stiffening of Cytoskeletal Components Regulates Cardiomyocyte Contractility

**DOI:** 10.1101/2025.02.28.640898

**Authors:** Mohammad Jafari, Sisir Datla, Elliot L. Elson, Brian E. Carlson, Tetsuro Wakatsuki, Farid Alisafaei

**Affiliations:** Department of Mechanical Engineering, New Jersey Institute of Technology, USA; Department of Mechanical Engineering and Materials Science, Washington University in St. Louis, USA; Department of Biochemistry and Molecular Biophysics, Washington University in St. Louis School of Medicine, USA; Department of Molecular and Integrative Physiology, University of Michigan, Ann Arbor, MI, USA; InvivoSciences, Inc., Madison, WI, USA

## Abstract

Cardiomyocytes continuously experience mechanical stimuli that regulate their contractile behavior and contribute to overall heart function. Despite the importance of mechanotransduction in cardiac physiology, the mechanisms by which cardiomyocytes integrate external mechanical cues, such as stretch and environmental stiffness, remain poorly understood. In this study, we present a combined theoretical and experimental framework to investigate how strain-induced cytoskeletal stiffening modulates cardiomyocyte contractility and force generation. Our study elucidates that regulating the mechanical tension cardiomyocytes experience in tissue—whether by modulating environmental stiffness, external stretching, or cardiac fibroblast activation—can effectively tune their contractility, with cytoskeletal strain stiffening playing a central role in this mechanotransductive response.

## Introduction

Cardiovascular disease remains the leading cause of death globally, contributing to a significant portion of mortality rates. In 2006, it accounted for approximately one in every 2.9 deaths in the United States.^1^ The rhythmic contractile activity of heart muscle cells, also called cardiac myocytes (CMs), is fundamental to maintaining blood circulation in mammals.^2^ Impairments in cardiomyocyte contractility are closely linked to cardiac dysfunction, and therapeutic strategies often focus on modulating CM contractile function to treat a failing heart.^3^

Single CMs contain the complete molecular machinery necessary for myocardial contraction,^4^ with contractile function driven by highly organized structures known as myofibrils.^5^ These myofibrils are composed of repeating units called sarcomeres, the basic contractile elements of muscle fibers.^6^ Each sarcomere consists primarily of actin and myosin protein filaments, whose interaction underlies muscular contraction.^7^ The coordinated activity of sarcomeric myosin, which hydrolyzes ATP to produce mechanical work,^8^ enables myosin filaments to slide along actin filaments, resulting in sarcomere shortening. This precisely regulated process drives the contraction of individual myofibrils, ultimately facilitating the contractile function of the heart.^2^

Cardiomyocytes, like many other cell types, possess the ability to sense and respond to mechanical cues from their environment, a process known as mechanosensing.^9^ The stiffness of the surrounding environment is a key parameter of the natural cardiac environment. In vitro studies have demonstrated that when cells are cultured on flat 2D substrates, increased substrate stiffness generally leads to elevated intracellular tension,^10,11^ which in turn affects cytoskeletal organization and promotes force generation.^12^ Similarly, in pathological conditions such as fibrosis, extracellular matrix (ECM) structural remodeling significantly alters the mechanical properties and stiffness of the myocardium, which profoundly impacts CM physiology and contractility,^13–18^ contributing to the progression of cardiac dysfunction.

In addition to environment stiffness, mechanical strain serves as another key regulator of cytoskeletal tension in cardiomyocytes. Strain level changes under various physiological and pathological conditions, including alterations in blood pressure, exercise, valvular diseases, and even the normal cardiac cycle.^19–21^ Despite the recognized influence of strain on myocardial function and its contraction force,^22^ the underlying mechanisms by which CMs sense and adapt to these dynamic mechanical cues remain poorly understood.

To investigate how cardiomyocytes respond to increased mechanical tension, whether originating from enhanced stiffness of the extracellular environment or from external stretching, we developed an integrated theoretical and experimental framework to examine the relationship between cardiomyocyte contractility and mechanical cues. Our findings demonstrated that cardiac force increases in response to both ECM stiffness and external mechanical stretching. Notably, the amplitude of contractile force, commonly referred to as twitch force, defined as the difference between peak and resting forces, also rises with increasing stretching. Our results suggest that strain-induced stiffening of the cytoskeleton components under tension plays a crucial role in the ability of cardiomyocytes to sense and respond to external mechanical stretching. This insight provides a potential explanation for how cytoskeletal mechanics contribute to the regulation of cardiac function and may play a role in the progression of cardiac dysfunction and heart failure.

## Results

### Cardiomyocyte contractility amplifies with environment stiffness

To investigate how environmental stiffness influences cardiac contractile force, we quantified changes in cardiomyocyte contractility *C*, reflecting the myosin-driven contractile response, and tensile stress *σ*, which represents the overall contraction capacity of the cardiomyocytes. The contractility of CMs arises from actomyosin activity, where myosin filaments slide along actin filaments, generating tension and driving contraction.^2^ During this process, distinct components of the cytoskeleton experience different mechanical stresses: structures such as microtubules undergo compression, whereas components like actin filaments are subjected to tension (Figure 1A). To explore how external stiffness affects cardiomyocyte contractility *C* and stress *σ*, we developed a chemo-mechanical model that treats the cytoskeleton as a continuum of representative volume elements (RVEs). Each RVE consisted of a contractile force-generating element, representing myosin,^23^ and two passive cytoskeletal elements (Figure 1B).^24^ One cytoskeletal element, arranged in parallel with the myosin, experienced compression during contraction, capturing the behavior of cytoskeletal components under compressive forces. The other element, connected in series with the myosin, underwent tension, reflecting cytoskeletal structures that experience tension during myosin contraction. This arrangement, as the CM, was connected to two passive elements on each side, representing the stiffness of the extracellular environment and allowing for the simulation of external mechanical constraints (Figure 1B).

**Figure 1.**
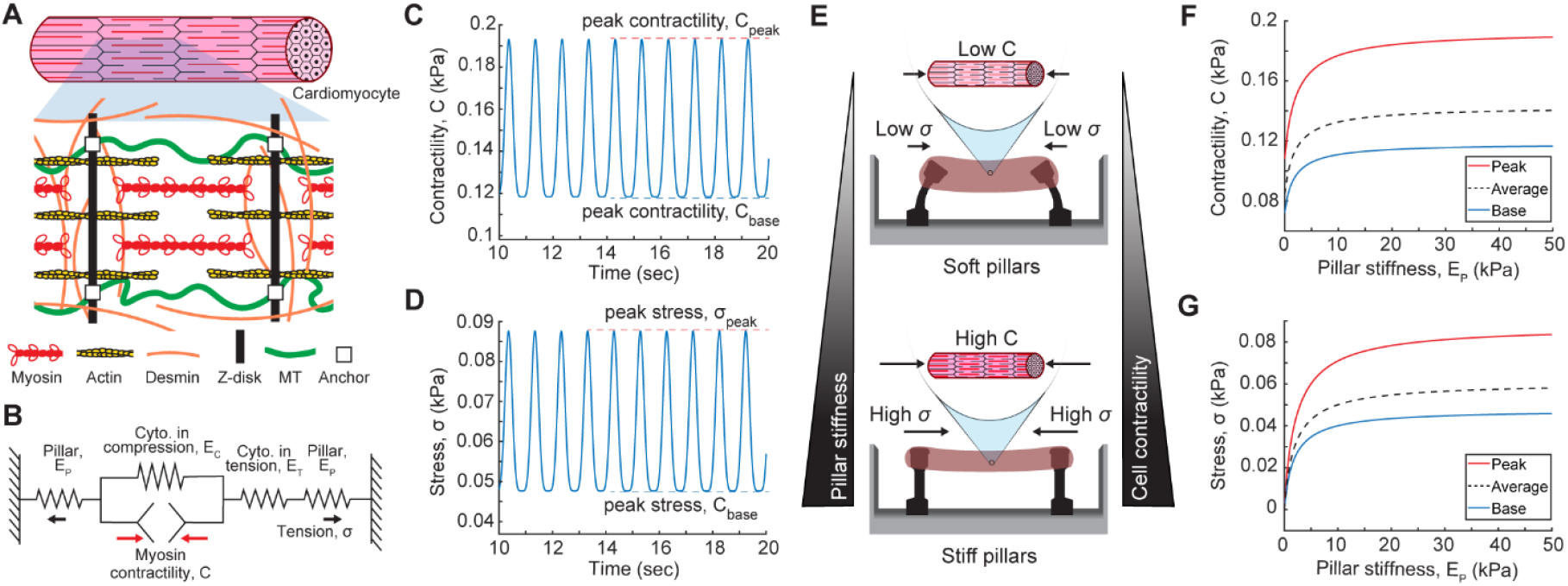
Cardiomyocyte contractility is enhanced by increased environment stiffness. (A) During cardiomyocyte contraction, cytoskeletal components such as actin experience tension, while other components, like microtubules, experience compression. (B) To study the effect of environment stiffness on cardiomyocyte behavior, we developed a chemo-mechanical cell model treating each cardiomyocyte as a representative volume element (RVE). Each RVE comprises a contractile, periodic force-generating element representing myosin contractility and two passive cytoskeletal components: one in parallel (cytoskeleton under compression, such as microtubules) and one in series (cytoskeleton under tension, such as actin) with the contractile element. (C) The myosin element generates a periodic contractility *C*, mimicking heartbeat dynamics with a base and peak cycle occurring every second. (D) A portion of this contractility is absorbed by cytoskeletal elements under compression, which resist myosin-driven contraction, while the rest is transmitted as tension, reflecting the stress cardiomyocytes generate (*σ*). (E) Schematics of a micropillar system with varying pillar stiffness. (F) The cell model predicted that stiffer pillars amplified the cardiomyocyte contractility *C* at both the base and peak magnitudes. (G) The enhanced cell contractility was also evidenced by the corresponding increases in the cell stress *σ* in baseline and amplitude.

The myosin element was modeled to generate a periodic contractile stress, *C*, simulating the cyclic nature of cardiac beating, with baseline and peak values, *C*_base_ and *C*peak, respectively, occurring every second (Figure 1C). In our model, we assumed a constant beating rate, consistent with experimental observations,^16^ which indicate that external mechanical cues, such as ECM stiffness, do not influence the frequency of myosin-generated contractile stress *C*. A portion of this contractile stress is absorbed by cytoskeletal elements under compression (e.g., microtubules),^25^ which resist myosin-driven contraction, while the remaining stress is transmitted as tensile stress, *σ*, reflecting the stress cardiomyocytes generate (Figure 1D).

Hereafter, we refer to *C*as cell contractility, as it represents the intrinsic tendency of the cell to contract, while we denote *σ* as cell stress, as it quantifies the actual stress generated by cell contractility. As described in the Supplementary Information, the central hypothesis of our model is that cell contractility *C* increases with increasing stress *σ*, reflecting the mechanosensitivity of cardiomyocytes. Increased mechanical stress is known to enhance actomyosin engagement through pathways involving stretch-activated ion channels, cytoskeletal remodeling, and calcium-dependent contractile regulation, allowing cardiomyocytes to adapt their force generation under varying mechanical loads.^26,27^ This mechanoresponsive behavior allows cardiomyocytes to adapt their contractility to maintain efficient force transmission and cardiac function under varying mechanical loads.

We first examined how changes in ECM stiffness influence cardiomyocyte contractility *C* and stress *σ*. To explore this, we conducted micropillar-based simulations in which CMs were anchored to pillars of varying stiffness *EP*(Figure 1E). The model predicted that increasing ECM stiffness led to an amplification of cardiomyocyte contractility *C* at both baseline and peak levels.

As shown in Figure 1F, stiffer pillars resulted in higher contractility, consistent with experimental findings demonstrating that increased substrate stiffness enhances CM contractility by generating sufficient intracellular tension, which is critical for regulating contractility and maintaining proper myofibril alignment.^28^

This increase in cell contractility was further reflected in the corresponding rise in contraction stress *σ* at both baseline and peak, as illustrated in Figure 1G. These predictions agree with previous experimental observations that reported increased contractile force in response to substrate stiffness in various samples, including human pluripotent stem cell (hPSC)-derived cardiomyocytes and neonatal rat cardiomyocytes,^16^ both cultured on substrates of varying stiffness. Similar results were observed in experiments involving rat cardiac myocytes anchored to elastic micropillars.^2^

The model also shows that pillar deflection decreased with increasing stiffness. As depicted in Figure S1A, both the peak and baseline displacements of the pillars *ε*_P_ decreased as pillar stiffness increased. Consistent with this trend, increased pillar stiffness also led to reduced cell shortening (Figure S1B). Considering the total length of a cardiomyocyte as (*ε*_C_+*ε*_T_), cell shortening was defined as the difference in cell length between baseline (resting state) and peak contraction. This prediction aligns with experimental observations from hPSC-derived cardiomyocytes, where relative cell shortening was significantly greater on softer polydimethylsiloxane (PDMS) substrates compared to those with higher stiffness.^9^

In our model, we assumed that myofibril integrity was maintained across all pillar stiffness conditions. This assumption reflects the critical role of structural connections within and between cardiomyocytes and ECM in ensuring coordinated contraction along a defined direction.^2^ Experimental evidence suggests that excessively high substrate stiffness can surpass a threshold where tension-induced myofibril rupture occurs, leading to a reduction in mechanical output and impaired contractility.^28^

Together, the model demonstrated that both cardiomyocyte contractility and contraction stress increased as pillar stiffness increased, reaching a plateau at higher stiffness levels. These findings highlighted the critical role of environmental stiffness in modulating cardiomyocyte contractility and stress. This behavior mirrored experimental observations, confirming the role of substrate stiffness in establishing intracellular tension to regulate contractility.

### Strain stiffening of the cytoskeleton regulates both the baseline and amplitude of cardiac contractility and stress

The cytoskeletal architecture is primarily composed of actin, microtubules, and intermediate filaments. Actin filaments and, to some extent, intermediate filaments experience tension (Figure 2A). Both actin networks^29,30^ and intermediate filaments^31–37^ exhibit nonlinear stiffening under tension. In our model, this strain-stiffening effect was incorporated by allowing the stiffness of the cytoskeletal element under tension *E*_T_ to increase in response to strain in that element (Figure 2B, S2).

**Figure 2.**
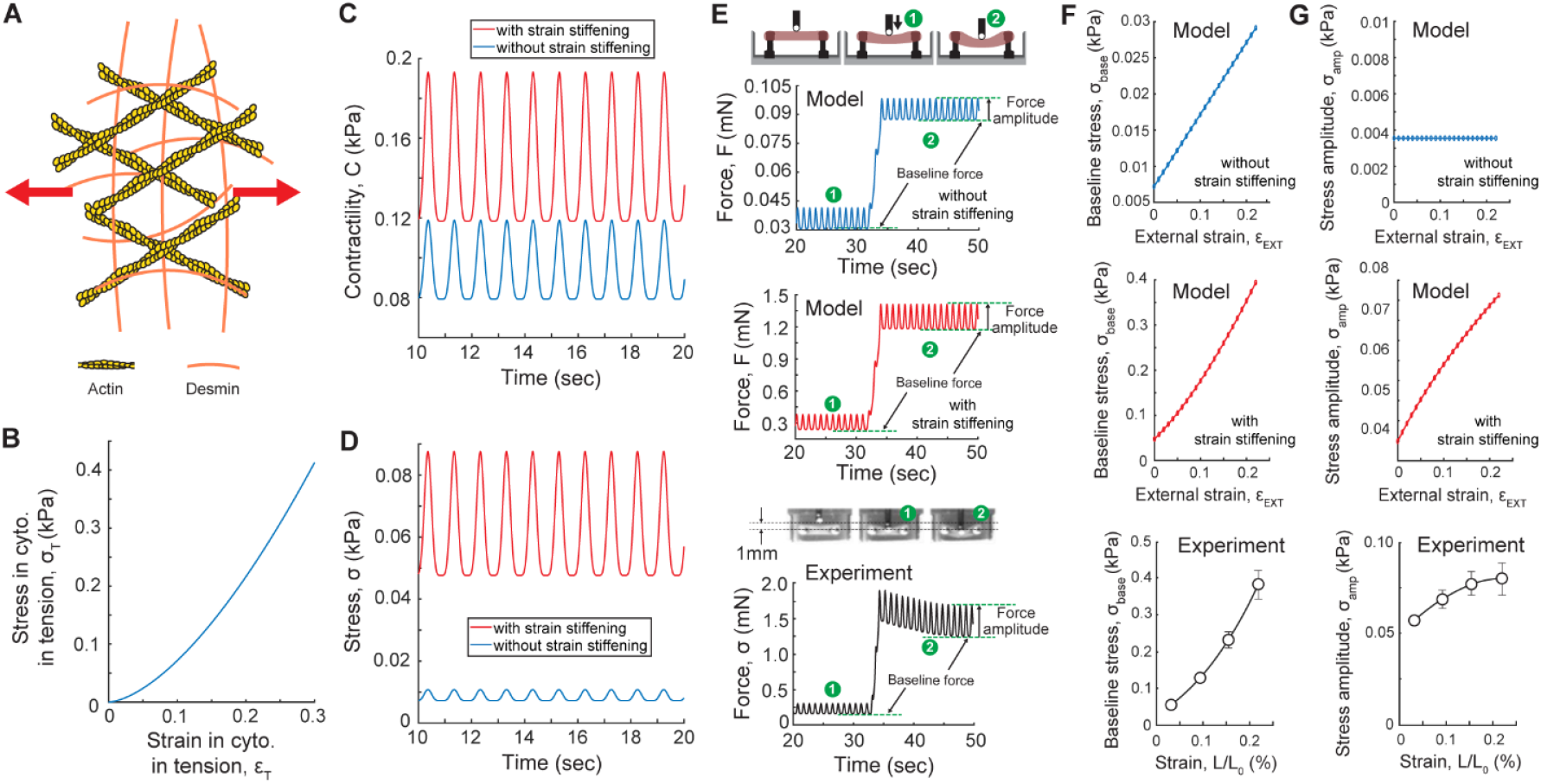
Strain-induced cytoskeleton stiffening enhances both the baseline and amplitude of cardiomyocyte stress. (A) As myosin contracts, actin filaments and, to some extent, intermediate filaments experience tension. (B) These components are known to exhibit a nonlinear stiffening response to tension, which was incorporated into our model by allowing the stiffness of the element under tension (*E*T) to increase in response to strain. This behavior is reflected in the increased slope of the stress-strain curve shown here. (C) The model predicted a significant increase in cell contractility when strain stiffening was incorporated, compared to simulations without this effect. (D) Stress measurements similarly reflected this enhancement, showing that strain stiffening substantially amplifies both *σ*_base_ and *σ*_amp_. (E) To investigate the effect of cytoskeletal strain-stiffening on cardiomyocyte force generation, we simulated the static stretching of cardiomyocytes. The model predicted that increasing the stretch level elevates the baseline stress *σ*_base_ in both scenarios; with and without the inclusion of strain stiffening. However, stress amplitude *σ*_amp_ increased only when strain stiffening was included, remaining unchanged without the strain stiffening effect. To determine which scenario more accurately reflects physiological behavior, model predictions were compared with experimental measurements of cardiac stress in engineered heart tissues (EHTs) subjected to two stretching levels. The experimental results aligned with the predictions incorporating strain stiffening. (F) Model predictions for baseline stress *σ*_base_ and (G) stress amplitude *σ*_amp_ in response to increasing external strain (*ε*_EXT_) were validated experimentally by measuring these stresses in EHTs subjected to four incremental strain levels (n = 6).

The model revealed that incorporating cytoskeletal strain stiffening significantly amplified both CM contractility *C* and stress *σ* (Figures 2C,D,S3), suggesting that this characteristic may play a crucial role in cardiomyocyte mechanotransduction. To experimentally test the effect of cytoskeletal strain-stiffening, we fabricated engineered heart tissues (EHTs), derived from human induced pluripotent stem cells (iPSCs), anchored to rigid posts. The tissue was stretched by pressing a round indenter tip vertically downward at its center, while simultaneously measuring the cyclic vertical force exerted by the tissue on the tip in the upward direction, representing the tissue’s cyclic contractility (Figure 2E). Additionally, we simulated our experimental setup by applying an external stretch (*ε*_EXT_) across the system and measuring the resulting cardiac stress (*σ*).

Model predictions showed that increasing external stretch elevated the baseline stress *σ*_base_ in both scenarios; with and without strain stiffening (Figure 2F), consistent with the Frank-Starling law of the heart, which states that cardiac muscle increases its contraction force in response to stretch within physiological limits.^22,28,38–41^ However, the model also revealed a critical distinction: while stress amplitude *σ*_amp_ increased with stretch in the presence of strain stiffening, it remained unchanged when strain stiffening was absent (Figure 2G). This raised a key question about which scenario more accurately reflects physiological behavior.

To address this, we measured *σ*_base_ and *σ*_amp_ in our experimental setup subjected to varying stretch levels. The experimental results closely matched the model predictions incorporating strain stiffening (Figures 2F,G), consistent with our previous observations showing that EHTs exhibit a stretch-dependent increase in stress amplitude *σ*_amp_,^22^ indicative of length-dependent activation mechanisms similar to those observed in the native heart.^42^

Both simulation and experimental results demonstrated that CMs stress amplitude *σ*_amp_ increased with external stretching but eventually exhibited saturation, as indicated by the diminishing slope in the stress-strain curve (Figure 2G). This finding aligns with earlier studies reporting a near-linear increase in force amplitude up to 15% strain, followed by a plateauing effect.^22^ In contrast, both experimental and theoretical results revealed that base stress *σ*_base_ and peak stress *σ*_peak_ showed a continuous, nonlinear increase with stretch (Figures 2F,S4), indicative of cytoskeletal strain stiffening.

These findings highlight the critical role of cytoskeletal strain stiffening in regulating cardiomyocyte mechanotransduction and force generation under external stretch. Incorporating strain-stiffening behavior into the model successfully captured key physiological responses, including the length-dependent activation of contraction force. When the cytoskeleton under tension was modeled as nonlinear (exhibiting strain-stiffening behavior), our simulations showed an increase in both baseline stress and stress amplitude with external stretch, aligning well with our experimental results. However, modeling the cytoskeleton under tension as a purely linear material resulted in an increase in baseline stress but no change in stress amplitude. These results show that strain-induced cytoskeletal remodeling is essential for proper cardiomyocyte function and plays a pivotal role in adapting cardiac force generation to mechanical stimuli.

### Increasing tissue tension elevates cardiomyocyte contractility

To further explore the impact of increased tension on cardiomyocyte contractility and force generation, we extended our model and experimental analysis to include cardiac fibroblasts. We fabricated engineered heart tissues (EHTs) using cardiomyocytes derived from human-induced pluripotent stem cells (iPSCs) combined with cardiac fibroblasts and collagen. To examine how cardiomyocytes respond to changes in tissue tension, we treated the tissues with transforming growth factor-beta (TGF-β), a known enhancer of fibroblast activation. This activation increases fibroblast contractility, generating higher tension within the tissue.

Our simulations showed an upregulation of myocyte contractility (*C*) and tissue stress (*σ*) following TGF-β treatment (Figures 3A, S5). Notably, when strain-stiffening of the cardiomyocyte cytoskeleton was included, both baseline stress (*σ*_base_, Figure 3B) and stress amplitude (*σ*_amp_, Figure 3C) increased with TGF-β treatment, and both also increased in response to tissue stretch (*ε*_EXT_). These findings aligned with our experiments, where stretching TGF-β-treated tissues revealed significant increases in both baseline stress and stress amplitude compared to untreated tissues, with both parameters further increasing under external stretch (Figures 3B and 3C). However, when strain stiffening was excluded from the model, although TGF-β treatment elevated both baseline stress and stress amplitude compared to untreated EHTs, the baseline response followed a linear trend, and the amplitude remained unchanged across different stretch levels (Figure 3D), which was inconsistent with our experimental results.

**Figure 3.**
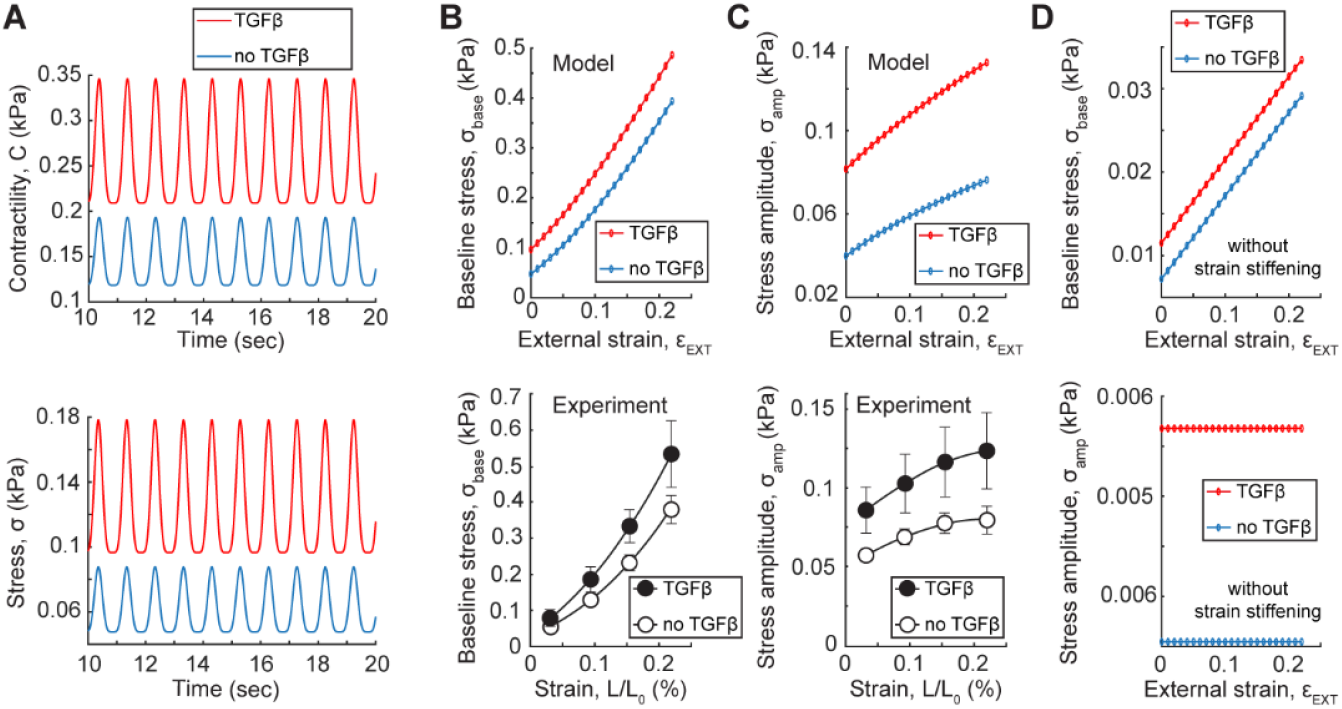
Increased cardiomyocyte contractility in response to elevated tissue tension. (A) Model simulations showing the upregulation of myocyte contractility (*C*) and tissue stress (*σ*) following TGF-β treatment. (B, C) Both baseline stress (*σ*_base_) and stress amplitude (*σ*_amp_) increased with TGF-β treatment, and both further increased under external stretch (*ε*_EXT_) when strain-stiffening was included in the model. These predictions were in agreement with our experiments, showing that both tissue baseline stress and stress amplitude increased with activation of cardiac fibroblasts upon TGF-β treatment, and both further increased under external stretch. (D) When strain-stiffening was excluded from the model, TGF-β treatment elevated baseline stress but did not affect stress amplitude across different stretch levels, which was inconsistent with experimental results.

Collectively, these results show that cardiomyocytes sense the tension within tissue and adjust their contractility accordingly. This was demonstrated by an increase in both the baseline and amplitude of their stresses in response to heightened tissue tension generated by activated fibroblasts.

### Regulation of myocyte contractility through fibroblast contractility

We next tested whether reducing tissue tension through deactivation of cardiac fibroblasts could decrease cardiomyocyte contractility. To this end, we reduced cardiac fibroblast contractility by inhibiting their non-muscle myosin using IVS201 treatment. IVS201, developed by InvivoSciences, is a repurposed therapeutic agent designed to target interstitial cardiac fibrosis, particularly in patients with heart failure with preserved ejection fraction (HFpEF). We hypothesize that inhibiting non-muscle myosin in fibroblasts will reduce their contractility, thereby lowering tissue tension. In response to this decreased tension, cardiomyocytes would then downregulate their muscle myosin.

Our simulations showed downregulation of myocyte contractility (*C*) and stress (*σ*) following the inhibition of fibroblast contractility (Figure 4A). Both baseline stress (*σ*_base_, Figure 4B) and stress amplitude (*σ*_amp_, Figure 4C) decreased with reduction of fibroblast contractility. These findings agreed with our experiments, where inhibiting non-muscle myosin in fibroblasts using IVS201 treatment led to a dose-dependent reduction in both baseline stress and stress amplitude.

**Figure 4.**
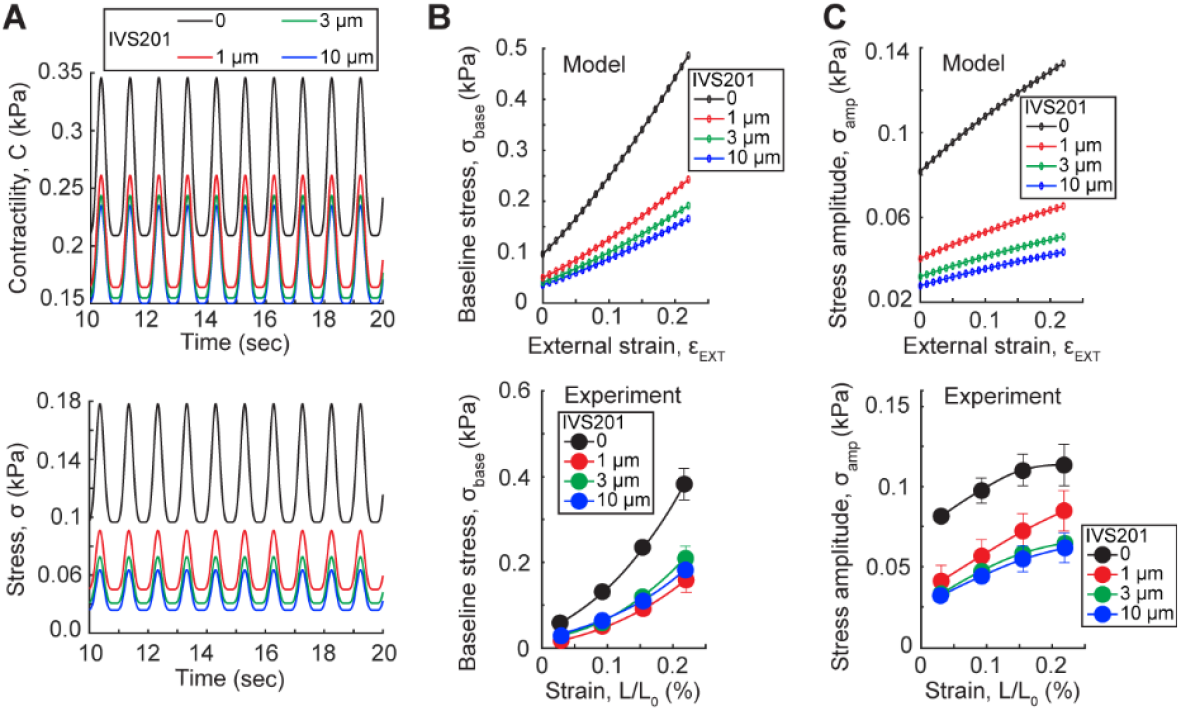
Reduction of fibroblast contractility lowers cardiomyocyte contractility and tissue stress. We studied theoretically and experimentally how the reduction of fibroblast contractility impacts myocyte contractility and overall tissue stress. (A) Model predictions showed a decrease in both baseline and amplitude of cardiomyocyte contractility *C*, with a more pronounced reduction in transmitted contractile stress *σ*, reflected in lower stress baseline and amplitude. (B) Predicted reductions in stress baseline *σ*_base_ and (C) amplitude σ_amp_ were validated experimentally by measuring these stresses in EHTs grown with TGF-β and treated with IVS201 for two weeks under different tissue stretch levels (*n* = 6).

Together, these findings revealed that reducing cardiac fibroblast contractility—without directly suppressing cardiomyocyte contractility—can expose cardiomyocytes to lower tension, which indirectly decreases both the baseline and amplitude of their contractility.

## Conclusions

Our study highlights the crucial role of nonlinear cytoskeletal strain stiffening in regulating cardiomyocyte contractility and mechanotransduction. By integrating theoretical modeling with experimental validation, we demonstrate that cytoskeletal strain stiffening enhances both the baseline and amplitude of contractile force in response to increased mechanical tension. Furthermore, we show that cardiomyocyte contractility can be modulated by controlling the mechanical tension they experience within the tissue. This can be achieved through adjustments in extracellular stiffness, external tissue stretching, or regulation of fibroblast-generated contractile forces. These findings provide fundamental insights into the mechanical regulation of cardiomyocytes and suggest potential strategies for tuning cardiac contractility in physiological and pathological conditions.

## References

1. Virani, S. S. et al. Heart Disease and Stroke Statistics—2020 Update: A Report From the American Heart Association. Circulation 141, e139–e596 ((2020).

2. Kajzar, A., Cesa, C. M., Kirchgeßner, N., Hoffmann, B. & Merkel, R. Toward Physiological Conditions for Cell Analyses: Forces of Heart Muscle Cells Suspended Between Elastic Micropillars. Biophysical Journal 94, 1854–1866 ((2008).

3. Dorn, G. W. & Molkentin, J. D. Manipulating Cardiac Contractility in Heart Failure. Circulation 109, 150–158 ((2004).

4. Louch, W. E., Sheehan, K. A. & Wolska, B. M. Methods in cardiomyocyte isolation, culture, and gene transfer. Journal of Molecular and Cellular Cardiology 51, 288–298 ((2011).

5. Ehler, E. Cardiac cytoarchitecture — why the “hardware” is important for heart function! Biochimica et Biophysica Acta (BBA) - Molecular Cell Research 1863, 1857–1863 ((2016).

6. de Tombe, P. P. & ter Keurs, H. E. D. J. The velocity of cardiac sarcomere shortening: mechanisms and implications. J Muscle Res Cell Motil 33, 431–437 ((2012).

7. Mansfield, P. J. & Neumann, D. A. Chapter 3 - Structure and Function of Skeletal Muscle. in Essentials of Kinesiology for the Physical Therapist Assistant (Third Edition) (eds. Mansfield, P. J. & Neumann, D. A. 34–49 (Mosby, St. Louis (MO), 2019). doi:10.1016/B978-0-323-54498-6.00003-5.

8. Cooper, G. M. Actin, Myosin, and Cell Movement. in The Cell: A Molecular Approach. 2nd edition (Sinauer Associates, 2000).

9. Körner, A., Mosqueira, M., Hecker, M. & Ullrich, N. D. Substrate Stiffness Influences Structural and Functional Remodeling in Induced Pluripotent Stem Cell-Derived Cardiomyocytes. Front. Physiol. 12, (2021).

10. Mih, J. D., Marinkovic, A., Liu, F., Sharif, A. S. & Tschumperlin, D. J. Matrix stiffness reverses the effect of actomyosin tension on cell proliferation. Journal of Cell Science 125, 5974–5983 ((2012).

11. Banerjee, S., Sknepnek, R. & Cristina Marchetti, M. Optimal shapes and stresses of adherent cells on patterned substrates. Soft Matter 10, 2424–2430 ((2014).

12. Chopra, A., Lin, V., McCollough, A., Atzet, S., Prestwich, G. D., Wechsler, A. S., Murray, M. E., Oake, S. A., Yasha Kresh, J. & Janmey, P. A. Reprogramming cardiomyocyte mechanosensing by crosstalk between integrins and hyaluronic acid receptors. Journal of Biomechanics 45, 824–831 ((2012).

13. Gaetani, R., Zizzi, E. A., Deriu, M. A., Morbiducci, U., Pesce, M. & Messina, E. When Stiffness Matters: Mechanosensing in Heart Development and Disease. Front. Cell Dev. Biol. 8, (2020).

14. Chopra, A., Tabdanov, E., Patel, H., Janmey, P. A. & Kresh, J. Y. Cardiac myocyte remodeling mediated by N-cadherin-dependent mechanosensing. American Journal of Physiology-Heart and Circulatory Physiology 300, H1252–H1266 ((2011).

15. Young, J. L., Kretchmer, K., Ondeck, M. G., Zambon, A. C. & Engler, A. J. Mechanosensitive Kinases Regulate Stiffness-Induced Cardiomyocyte Maturation. Sci Rep 4, 6425 ((2014).

16. Hazeltine, L. B., Simmons, C. S., Salick, M. R., Lian, X., Badur, M. G., Han, W., Delgado, S. M., Wakatsuki, T., Crone, W. C., Pruitt, B. L. & Palecek, S. P. Effects of Substrate Mechanics on Contractility of Cardiomyocytes Generated from Human Pluripotent Stem Cells. International Journal of Cell Biology 2012, 508294 ((2012).

17. Hersch, N., Wolters, B., Dreissen, G., Springer, R., Kirchgeßner, N., Merkel, R. & Hoffmann, B. The constant beat: cardiomyocytes adapt their forces by equal contraction upon environmental stiffening. Biology Open 2, 351–361 ((2013).

18. Ward, M. & Iskratsch, T. Mix and (mis-)match – The mechanosensing machinery in the changing environment of the developing, healthy adult and diseased heart. Biochimica et Biophysica Acta (BBA) - Molecular Cell Research 1867, 118436 ((2020).

19. Rysä, J., Tokola, H. & Ruskoaho, H. Mechanical stretch induced transcriptomic profiles in cardiac myocytes. Sci Rep 8, 4733 ((2018).

20. Brady, B., King, G., Murphy, R. T. & Walsh, D. Myocardial strain: a clinical review. Ir J Med Sci 1–8 (2022) doi:10.1007/s11845-022-03210-8.

21. Neves, J. S., Leite-Moreira, A. M., Neiva-Sousa, M., Almeida-Coelho, J., Castro-Ferreira, R. & Leite-Moreira, A. F. Acute Myocardial Response to Stretch: What We (don’t) Know. Front Physiol 6, 408 ((2016).

22. Asnes, C. F., Marquez, J. P., Elson, E. L. & Wakatsuki, T. Reconstitution of the Frank-Starling Mechanism in Engineered Heart Tissues. Biophysical Journal 91, 1800–1810 ((2006).

23. Shenoy, V. B., Wang, H. & Wang, X. A chemo-mechanical free-energy-based approach to model durotaxis and extracellular stiffness-dependent contraction and polarization of cells. Interface Focus 6, 20150067 ((2016).

24. Alisafaei, F., Jokhun, D. S., Shivashankar, G. V. & Shenoy, V. B. Regulation of nuclear architecture, mechanics, and nucleocytoplasmic shuttling of epigenetic factors by cell geometric constraints. PNAS 116, 13200–13209 ((2019).

25. Robison, P., Caporizzo, M. A., Ahmadzadeh, H., Bogush, A. I., Chen, C. Y., Margulies, K. B., Shenoy, V. B. & Prosser, B. L. Detyrosinated microtubules buckle and bear load in contracting cardiomyocytes. Science 352, aaf0659 ((2016).

26. Kane, C., Couch, L. & Terracciano, C. M. N. Excitation–contraction coupling of human induced pluripotent stem cell-derived cardiomyocytes. Front. Cell Dev. Biol. 3, (2015).

27. Bers, D. M. Cardiac excitation–contraction coupling. Nature 415, 198–205 ((2002).

28. Ribeiro, A. J. S., Ang, Y.-S., Fu, J.-D., Rivas, R. N., Mohamed, T. M. A., Higgs, G. C., Srivastava, D. & Pruitt, B. L. Contractility of single cardiomyocytes differentiated from pluripotent stem cells depends on physiological shape and substrate stiffness. Proceedings of the National Academy of Sciences 112, 12705–12710 ((2015).

29. Xu, J., Tseng, Y. & Wirtz, D. Strain Hardening of Actin Filament Networks: REGULATION BY THE DYNAMIC CROSS-LINKING PROTEIN α-ACTININ*. Journal of Biological Chemistry 275, 35886–35892 ((2000).

30. Åström, J. A., Kumar, P. B. S., Vattulainen, I. & Karttunen, M. Strain hardening, avalanches, and strain softening in dense cross-linked actin networks. Phys. Rev. E 77, 051913 ((2008).

31. Ma, L., Xu, J., Coulombe, P. A. & Wirtz, D. Keratin Filament Suspensions Show Unique Micromechanical Properties *. Journal of Biological Chemistry 274, 19145–19151 ((1999).

32. Janmey, P. A., Euteneuer, U., Traub, P. & Schliwa, M. Viscoelastic properties of vimentin compared with other filamentous biopolymer networks. Journal of Cell Biology 113, 155–160 ((1991).

33. Ackbarow, T. & Buehler, M. J. Superelasticity, energy dissipation and strain hardening of vimentin coiled-coil intermediate filaments: atomistic and continuum studies. J Mater Sci 42, 8771–8787 ((2007).

34. Charrier, E. E. & Janmey, P. A. Chapter Two - Mechanical Properties of Intermediate Filament Proteins. in Methods in Enzymology (eds. Omary, M.B. & Liem, R.K.H.) vol. 568 35–57 (Academic Press, 2016).

35. Lin, Y.-C., Yao, N. Y., Broedersz, C. P., Herrmann, H., MacKintosh, F. C. & Weitz, D. A. Origins of Elasticity in Intermediate Filament Networks. Phys. Rev. Lett. 104, 058101 ((2010).

36. Pegoraro, A. F., Janmey, P. & Weitz, D. A. Mechanical Properties of the Cytoskeleton and Cells. Cold Spring Harb Perspect Biol 9, a022038 ((2017).

37. Alisafaei, F., Mandal, K., Saldanha, R., Swoger, M., Yang, H., Shi, X., Guo, M., Hehnly, H., Castañeda, C. A., Janmey, P. A., Patteson, A. E. & Shenoy, V. B. Vimentin is a key regulator of cell mechanosensing through opposite actions on actomyosin and microtubule networks. Commun Biol 7, 1– 13 ((2024).

38. Pfeffer, M. A. & Braunwald, E. Ventricular remodeling after myocardial infarction. Experimental observations and clinical implications. Circulation 81, 1161–1172 ((1990).

39. Sarnoff, S. J. & Berglund, E. Ventricular Function. Circulation 9, 706–718 ((1954).

40. Holubarsch, C., Ruf, T., Goldstein, D. J., Ashton, R. C., Nickl, W., Pieske, B., Pioch, K., Lüdemann, J., Wiesner, S., Hasenfuss, G., Posival, H., Just, H. & Burkhoff, D. Existence of the Frank-Starling Mechanism in the Failing Human Heart. Circulation 94, 683–689 ((1996).

41. ter Keurs, H. E., Rijnsburger, W. H., van Heuningen, R. & Nagelsmit, M. J. Tension development and sarcomere length in rat cardiac trabeculae. Evidence of length-dependent activation. Circulation Research 46, 703–714 ((1980).

42. Tavi, P., Han, C. & Weckström, M. Mechanisms of Stretch-Induced Changes in [Ca2+]i in Rat Atrial Myocytes. Circulation Research 83, 1165–1177 ((1998).

