## Supplementary Information for "Tension-Induced Stiffening of Cytoskeletal Components Regulates Cardiomyocyte Contractility"

### 1D chemomechanical cell model

To elucidate how cardiomyocytes (CMs) respond to external mechanical stimuli, we developed a one-dimensional (1D) chemo-mechanical cell model. This simplified framework integrates both active and passive mechanical elements to simulate the dynamic interaction between CM components and external forces. The model consists of a single active contractile element, representing myosin-driven contractility, and two passive elements, representing cytoskeletal components subjected to either compression or tension during contraction. When the myosin contracts, certain cytoskeletal components experience tension, while others undergo compression (Figure 1A). To capture this, one passive element is connected in parallel with the active myosin element to represent cytoskeletal components under compression, while another passive element is connected in series to represent components under tension. To mimic the experimental configuration where cells are anchored between two pillars, the entire cell model, including both active and passive elements, is connected in series with two additional elements, which simulate the mechanical stiffness of the anchoring pillars (Figure 1B).

### 1. Cytoskeleton in compression

In this model, cytoskeletal components that experience compression during myosin contraction are represented by a linear elastic element connected in parallel with the contractile myosin element. For simplicity, we considered this element as a linear spring with a stiffness of $E_c$. The compressive stress developed in this element is given by $E_c \varepsilon_c$, where $\varepsilon_c$ represents the strain within the compressed cytoskeletal element and is equal to the myosin strain as this element is connected in parallel with the myosin.

### 2. Myosin

The core of the model is designed based on the observation that myosin contractility rises in response to cytoskeletal tension $\sigma$,[1,2] a phenomenon well-documented across various cell types.[3–6] This tension can originate from either myosin-driven intrinsic contractility or external mechanical forces. We start with deriving the equation for the total free energy $W$ of the myosin and the parallel element, which includes the following terms:

$$W(C,\varepsilon_{\mathrm{C}}) = C\varepsilon_{\mathrm{C}} + \frac{\beta}{2}(C - C_0)^2 - \alpha\int_0^{\rho}\sigma\, dC + \frac{1}{2}E_{\mathrm{C}}\,\varepsilon_{\mathrm{C}}^{\,2} - \int_0^{\varepsilon_{\mathrm{C}}}\sigma\, d\varepsilon_{\mathrm{C}} \tag{S1}$$

In this equation, the first term represents the mechanical work done by the myosin element, where $C$ denotes contractile stress, and $\varepsilon_{\mathrm{C}}$ is the strain in the myosin element (equal to the strain of the cytoskeletal element under compression). The second and third terms account for the chemical energy, with the second term capturing the rise in free energy due to deviations from the baseline contractility $C_0$ (contractility in the absence of external tension), and the third term representing the energy reduction resulting from the recruitment of molecular motors through tension-dependent signaling pathways, such as $Ca^{2+}$ activation.[1,2] The parameters $\beta$ and $\alpha$ regulate the chemo-mechanical feedback mechanisms. Specifically, $\beta$ controls the resistance against increased contractility in response to tension, while $\alpha$ modulates the activation of tension-dependent pathways. The fourth term captures the elastic energy stored in the cytoskeletal components under compression, and the fifth term reflects the mechanical work done by the cytoskeletal tension acting on the contractile machinery.

The baseline myosin-generated contractility was modeled as a periodic function to reflect the cyclic beating nature of cardiomyocytes. To achieve this, the baseline contractility $C_0$ was defined as:

$$C_0 = C_{0_{\mathrm{base}}} + C_{0_{\mathrm{amp}}} = C_{0_{\mathrm{base}}} + amplitude * |\sin \pi t|^n \tag{S2}$$

where $C_{0_{\mathrm{base}}}$ and $amplitude$ are constants, $t$ represents time (in seconds), and $n$ controls the sharpness of the contractile peaks. For all simulations, $n$ was set to 5 to replicate the sharp contraction-relaxation dynamics characteristic of cardiac cycles. Parameter values are summarized in Table S1.

To determine how myosin strain and contractility change with cytoskeletal tension, we minimized the total free energy. This requires setting the derivatives of $W$ with respect to $C$ and $\varepsilon_{\mathrm{C}}$ to zero:

$$\frac{\partial \mathrm{W}(C,\varepsilon_{\mathrm{C}})}{\partial \varepsilon_C} = C + E_{\mathrm{C}}\,\varepsilon_{\mathrm{C}} - \sigma = 0 \tag{S3}$$

$$\frac{\partial \mathrm{W}(C,\varepsilon_{\mathrm{C}})}{\partial C} = \varepsilon_{\mathrm{C}} + \beta(C - C_0) - \alpha\sigma = 0 \tag{S4}$$

From equation (S3), contractility can be expressed as:

$$C = \sigma - E_{\mathrm{C}}\,\varepsilon_{\mathrm{C}} \tag{S5}$$

This relation illustrates how myosin-generated contractility is used for compacting the cytoskeleton components in compression, while the remaining stress is transmitted as tension $\sigma$ within the cytoskeleton components in tension. In the absence of external stretching, $\varepsilon_C$ is negative due to intrinsic myosin contraction. Substituting the expression for $\sigma$ from equation (S4) into equation (S5) gives the following relation between contractility and myosin strain at a given time:

$$C = \frac{E_C\,\alpha - 1}{\beta - \alpha}\varepsilon_C + \frac{\beta}{\beta - \alpha}C_0 \tag{S6}$$

Furthermore, replacing $\varepsilon_C$ from equation (S5) into equation (S6) the previous equation gives the following expression for the relationship between contractility and cytoskeletal tension:

$$C = \frac{E_C\,\alpha - 1}{E_C\,\beta - 1}\sigma + \frac{E_C\,\beta}{E_C\,\beta - 1}C_0 \tag{S7}$$

This equation illustrates how tension transmitted through the cytoskeleton enhances contractile activity, consistent with mechanotransduction observations in cardiomyocytes. The model predicts that contractility varies within a specific range depending on mechanical constraints. Minimum contractility occurs when the cell contracts freely without resistance ($\sigma = 0$), while maximum contractility is achieved when contraction is fully restricted ($\varepsilon_C = 0$). Therefore, the range of contractility in the absence of external stretching can be described by:

$$\frac{E_C\beta C_0}{E_C\beta - 1} < C < \frac{\beta C_0}{\beta - 1} \tag{S8}$$

### 3. Cytoskeleton in tension

The cytoskeletal components subjected to tension are represented by an element connected in series with the contractile element. As outlined in the main text, myosin contraction generates forces that pull on actin and intermediate filaments. When subjected to tension, both actin filament networks[7,8] and intermediate filaments[9–14] exhibit nonlinear stiffening.

To accurately capture this behavior, the tensile element is modeled as a nonlinear elastic material, reflecting the strain-stiffening properties of the cytoskeleton under tension. The stiffness of this element $E_T$ is defined as a strain-dependent function:

$$E_T = E_{T_0} + l\varepsilon_T{}^m \tag{S9}$$

Where $E_{T_0}$ denotes the initial stiffness of this element, $\varepsilon_T$ is the strain generated in this element, and $l$ and $m$ are material-specific parameters that define the degree of strain stiffening. The corresponding stress $\sigma$, which reflects the stress transmitted through the cytoskeletal network, is determined by integrating the stiffness over the range of applied strain:

$$\sigma = \int_0^{\varepsilon_T} E_T\, d\varepsilon_T = \int_0^{\varepsilon_T} (E_{T_0} + l\varepsilon_T{}^m)\, d\varepsilon_T = E_{T_0}\varepsilon_T + \frac{l}{m+1}\varepsilon_T{}^{m+1} \tag{S10}$$

This expression captures the nonlinear increase in stress $\sigma$ with strain $\varepsilon_T$, modeling the strain-stiffening behavior characteristic of cytoskeletal filaments.

### 4. Coupling of the pillars and cell model

In the model, the cytoskeletal tension generated by actomyosin activity is transmitted to the anchoring pillars at both ends. The pillars are modeled as linear elastic elements with a known stiffness $E_{\mathrm{P}}$, allowing the transmitted stress $\sigma$ to be described as:

$$\sigma = E_P \varepsilon_{\mathrm{P}} \tag{S11}$$

To complete the system of equations, boundary conditions must be applied, reflecting the pillar-based experimental configuration where both ends of the system are fixed. Under this condition, the sum of the strains across all elements connected in series must be zero:

$$\varepsilon_C + \varepsilon_{\mathrm{T}} + 2\varepsilon_{\mathrm{P}} = 0 \tag{S12}$$

Note that, for the stretching experiments, where cells are subjected to external mechanical stretching, the boundary condition is modified to incorporate the externally applied strain $\varepsilon_{\mathrm{EXT}}$:

$$\varepsilon_C + \varepsilon_{\mathrm{T}} + 2\varepsilon_{\mathrm{P}} = \varepsilon_{\mathrm{EXT}} \tag{S13}$$

By integrating this boundary condition into the system, the full set of unknown variables $\sigma$, $C$, $\varepsilon_{\mathrm{C}}$, $\varepsilon_{\mathrm{T}}$, and $\varepsilon_{\mathrm{P}}$ can be determined by solving the system of equations composed of Equations (S3), (S4), (S10), (S11), and the corresponding boundary condition (either Equation (S12) or (S13), depending on the experimental context.

### 5. Stability criterion

To ensure the physical validity of the model, the myosin contractility $C$ must remain positive at all times, reflecting the nature of myosin-generated contractility, which exerts a pulling force on actin filaments rather than a compressive one. For this condition to hold, the coefficients in Equation (S6) must be positive:

$$\begin{gathered} \frac{E_C \alpha - 1}{\beta - \alpha} > 0 \\ \frac{\beta}{\beta - \alpha} > 0 \end{gathered} \tag{S14}$$

Additionally, $\beta$ and $\alpha$ must not be equal, as their equality would lead to a singularity in Equation (S6), resulting in unbounded values for contractility $C$ and, consequently, for stress $\sigma$, which are not physically feasible within the constraints of biological systems. Furthermore, the parameter $E_{\mathrm{C}}$, representing the stiffness of the cytoskeletal elements under compression, must always remain positive, as a negative stiffness value is not physically meaningful. Together, these conditions impose the following stability criterion for the system:

$$\frac{1}{E_{\mathrm{C}}} < \alpha < \beta \tag{S15}$$

### SI Figures

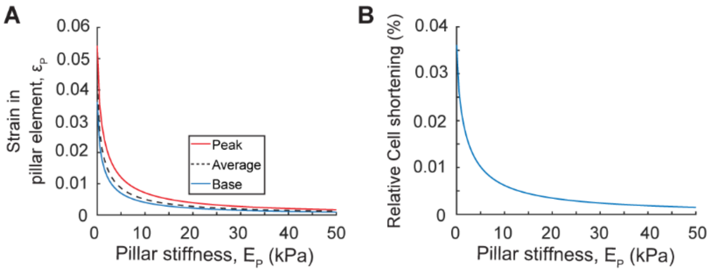


Figure S1. Model predictions for the effect of pillar stiffness $E_P$ on pillar strain and cell shortening. (A) Increasing pillar stiffness $E_P$ resulted in a decrease in pillar strain $\varepsilon_P$. (B) Increased pillar stiffness reduced cell shortening, defined as the difference between peak and baseline cell strain magnitudes $(\varepsilon_C + \varepsilon_T)$.

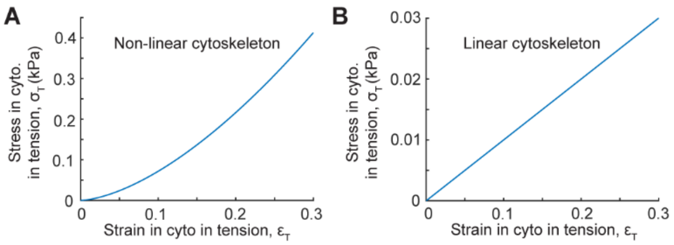


Figure S2. Strain stiffening in the cytoskeleton components under tension. During cell contraction, part of the cytoskeleton components is pulled. This tension leads to a significant localized increase in stiffness. In our chemomechanical cell model, we assumed that cytoskeletal components experiencing tension undergo stiffening in response to this tension. (A) The strain-stiffening behavior of the cytoskeleton in our model was reflected as an increased slope in the stress-strain curve in the cytoskeleton element under tension. The slope of the curve represents the element stiffness $E_{\mathrm{T}}$, starting from an initial low slope of $E_{\mathrm{T0}}$ and increasing nonlinearly with strain in this element $\varepsilon_{\mathrm{T}}$. (B) The stress-strain curve of cytoskeleton element under tension without strain stiffening in our model, exhibiting a fixed low slope as stiffness $E_{\mathrm{T0}}$ independent of the stretching level.

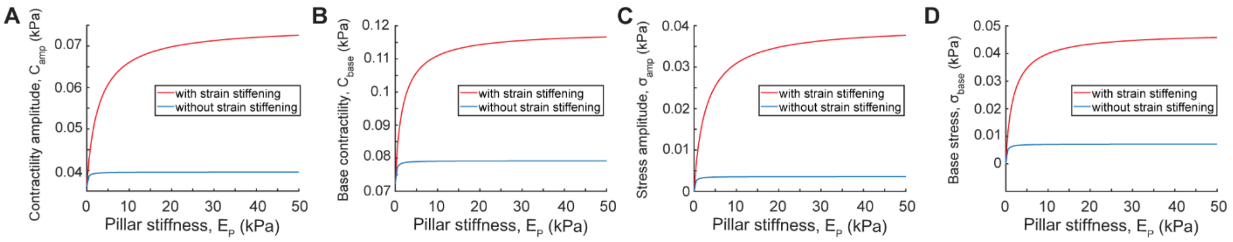


Figure S3. Strain-induced cytoskeleton stiffening enhances both contractility and stress baseline and amplitude. To investigate the impact of cytoskeletal strain stiffening on cell contractility and transmitted stresses, the chemomechanical cell model was used to quantify the baseline and amplitude of CM contractility in the micropillar experiment with and without strain stiffening effect. (A) Under the same pillar stiffness, the model predicted a significant increase in the amplitude of CM contractility $C_{\mathrm{amp}}$ due to strain stiffening in cytoskeletal components under tension. (B) Base cell contractility $C_{\mathrm{base}}$ followed a similar trend as the amplitude, with strain stiffening leading to a substantial increase compared to the case without strain stiffening. (C) Stress amplitude $\sigma_{\mathrm{amp}}$ and (D) baseline stress $\sigma_{\mathrm{base}}$ also reflected the same effect of strain stiffening.

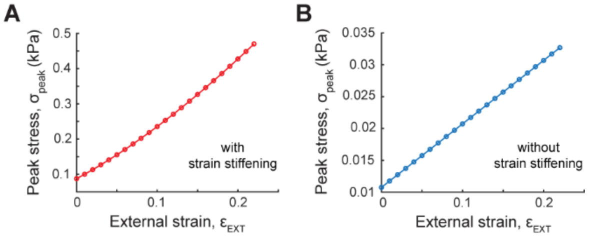


Figure S4. Model prediction for the effect of external stretching on the peak stress. (A) The model predicted a nonlinear increase in peak stress $\sigma_{\mathrm{peak}}$, defined as the sum of baseline stress $\sigma_{\mathrm{base}}$ and stress amplitude $\sigma_{\mathrm{amp}}$, in response to external stretching $\varepsilon_{\mathrm{EXT}}$. This nonlinearity comes from the strain stiffening of the cytoskeleton. (B) In the absence of strain stiffening, the peak stress exhibited a linear increase, mirroring the behavior of baseline stress $\sigma_{\mathrm{base}}$ (Figure 2F).

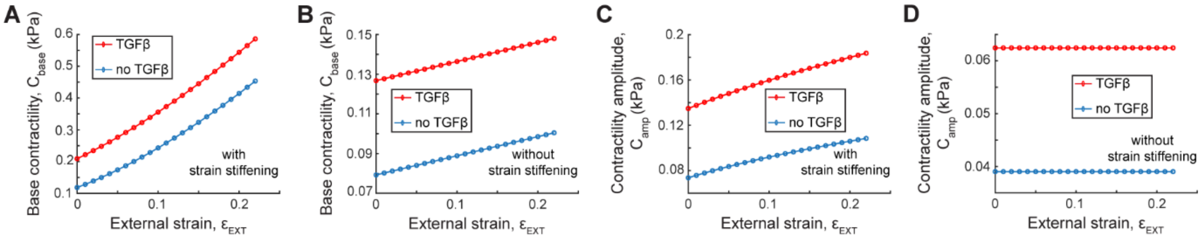


Figure S5. Model predictions for cardiomyocyte contractility in tissues grown with TGF-β. (A) Cardiomyocyte baseline contractility $C_{\mathrm{base}}$ increased in tissues treated with TGF-β, while maintaining a nonlinear relationship with external stretching. (B) In the absence of strain stiffening, model predictions showed that $C_{\mathrm{base}}$ increased with TGF-β; however, similar to untreated tissues, it exhibited a linear increase in response to external stretch. (C) The model also predicted an elevation in contractility amplitude $C_{\mathrm{amp}}$ with TGF-β treatment, preserving the nonlinear relationship with stretch. (D) Without strain stiffening, $C_{\mathrm{amp}}$ increased with TGF-β but remained unchanged in response to external stretch.

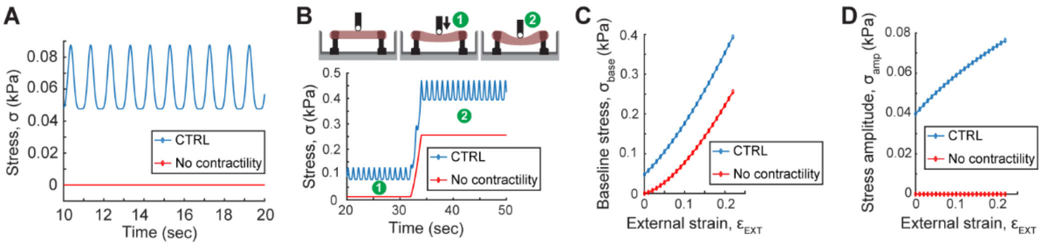


Figure S6. Model predictions for cardiomyocyte contractility abruption. (A) In the absence of cardiomyocyte contractility $C = 0$, tissues attached to pillars exhibited no stress $\sigma$ when unstretched, regardless of pillar stiffness. (B-D) Model simulations predicted that applying external stretch $\varepsilon_{\mathrm{EXT}}$ to these non-contractile tissues led to a nonlinear increase in baseline stress $\sigma_{\mathrm{base}}$, reflecting the strain-stiffening behavior of the cytoskeletal components under tension.

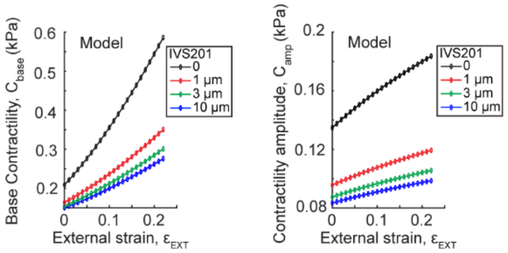


Figure S7. Model predictions for CM contractility when treating with IVS201, demonstrating a decline in both base and amplitude.

### Tables

Table S1 Parameters for 1D chemomechanical cell model.

| Parameter | Definition | Unit | Value |
|---|---|---|---|
| ${E_T}^0$ | Initial stiffness of cytoskeletal components in tension | kPa | 0.1 |
| $E_C$ | Stiffness of cytoskeletal components in compression | kPa | 1 |
| $\beta$ | Chemical stiffness parameter | kPa | 32 |
| $\alpha$ | Feedback parameter | kPa | 31 |
| ${C^0}_{\mathrm{base}}$ | CM contractility baseline in the absence of external stress | kPa | 0.07 |
| ${C^0}_{\mathrm{amplitude}}$ | CM contractility amplitude in the absence of external stress | kPa | 0.035 |
| $l$ | Strain stiffening parameter for the cytoskeleton in tension | | 45 |
| $m$ | Strain stiffening parameter for the cytoskeleton in tension | | 0.6 |

**References**


1. Bers, D. M. Cardiac excitation–contraction coupling. *Nature* **415**, 198–205 (2002).
2. Kane, C., Couch, L. & Terracciano, C. M. N. Excitation–contraction coupling of human induced pluripotent stem cell-derived cardiomyocytes. *Front. Cell Dev. Biol.* **3**, (2015).
3. Shenoy, V. B., Wang, H. & Wang, X. A chemo-mechanical free-energy-based approach to model durotaxis and extracellular stiffness-dependent contraction and polarization of cells. *Interface Focus* **6**, 20150067 (2016).
4. Lee, H., Alisafaei, F., Adebawale, K., Chang, J., Shenoy, V. B. & Chaudhuri, O. The nuclear piston activates mechanosensitive ion channels to generate cell migration paths in confining microenvironments. *Science Advances* **7**, eabd4058 (2021).
5. Matthews, B. D., Thodeti, C. K., Tytell, J. D., Mammoto, A., Overby, D. R. & Ingber, D. E. Ultra-rapid activation of TRPV4 ion channels by mechanical forces applied to cell surface β1 integrins. *Integrative Biology* **2**, 435–442 (2010).
6. Kobayashi, T. & Sokabe, M. Sensing substrate rigidity by mechanosensitive ion channels with stress fibers and focal adhesions. *Current Opinion in Cell Biology* **22**, 669–676 (2010).
7. Xu, J., Tseng, Y. & Wirtz, D. Strain Hardening of Actin Filament Networks: REGULATION BY THE DYNAMIC CROSS-LINKING PROTEIN α-ACTININ*. *Journal of Biological Chemistry* **275**, 35886–35892 (2000).
8. Åström, J. A., Kumar, P. B. S., Vattulainen, I. & Karttunen, M. Strain hardening, avalanches, and strain softening in dense cross-linked actin networks. *Phys. Rev. E* **77**, 051913 (2008).
9. Ma, L., Xu, J., Coulombe, P. A. & Wirtz, D. Keratin Filament Suspensions Show Unique Micromechanical Properties *. *Journal of Biological Chemistry* **274**, 19145–19151 (1999).
10. Janmey, P. A., Euteneuer, U., Traub, P. & Schliwa, M. Viscoelastic properties of vimentin compared with other filamentous biopolymer networks. *Journal of Cell Biology* **113**, 155–160 (1991).
11. Ackbarow, T. & Buehler, M. J. Superelasticity, energy dissipation and strain hardening of vimentin coiled-coil intermediate filaments: atomistic and continuum studies. *J Mater Sci* **42**, 8771–8787 (2007).
12. Charrier, E. E. & Janmey, P. A. Chapter Two - Mechanical Properties of Intermediate Filament Proteins. in *Methods in Enzymology* (eds. Omary, M. B. & Liem, R. K. H.) vol. 568 35–57 (Academic Press, 2016).
13. Lin, Y.-C., Yao, N. Y., Broedersz, C. P., Herrmann, H., MacKintosh, F. C. & Weitz, D. A. Origins of Elasticity in Intermediate Filament Networks. *Phys. Rev. Lett.* **104**, 058101 (2010).
14. Pegoraro, A. F., Janmey, P. & Weitz, D. A. Mechanical Properties of the Cytoskeleton and Cells. *Cold Spring Harb Perspect Biol* **9**, a022038 (2017).
